# Detection of Discordant Peptide Quantities in Shotgun Proteomics Data by Peptide Correlation Analysis (PeCorA)

**DOI:** 10.1101/2020.08.21.261818

**Authors:** Jesse G. Meyer

**Affiliations:** Department of Biochemistry, Medical College of Wisconsin, 8701 Watertown Plank Rd. Milwaukee, WI 53226

## Abstract

Shotgun proteomics techniques infer the presence and quantity of proteins using peptide proxies, which are produced by cleavage of all isolated protein by a protease. Most protein quantitation strategies assume that multiple peptides derived from a protein will behave quantitatively similar across treatment groups, but this assumption may be false for biological or technical reasons. Here, I describe a strategy called peptide correlation analysis (PeCorA) that detects quantitative disagreements between peptides mapped to the same protein. Simple linear models are used to assess whether the slope of a peptide’s change across treatment groups differs from the slope of all other peptides assigned to the same protein. Reanalysis of proteomic data from primary mouse microglia with PeCorA revealed that about 15% of proteins contain one discordant peptide. Inspection of the discordant peptides shows utility of PeCorA for direct and indirect detection of regulated PTMs, and also for discovery of poorly quantified peptides that should be excluded. PeCorA can be applied to an arbitrary list of quantified peptides, and is freely available as a script written in R.

## Introduction

Shotgun proteomics relies on the inference of protein identity and quantity from peptide pieces. Proteins are regulated before mRNA translation through alternative splicing of mRNA transcripts, and after translation by post-translational modifications (PTMs); thus, different subsections of the whole protein sequence might be influenced differently by biological perturbations. Such fine-grained protein regulation may be responsible for the diversity of function from our genome, and enables precise control of metabolism and gene expression. However, proteomic analysis is almost entirely based on combining multiple peptide quantities unique to a protein into one quantity^1,2^.

Although the magnitude of peptide abundance will differ due to different detection sensitivity^3,4^, most protein quantitation strategies assume that multiple peptides derived from a protein will behave quantitatively similar across treatment groups. For example, if a treatment causes a protein to increase, we assume that all peptides mapped to that protein will increase (**Figure 1A**). Because each protein encoded by a gene exists as a population of unique states, called proteoforms^5^, there are many biological reasons that multiple peptides from a protein may disagree quantitatively. For example, a change in the proportions of one gene’s proteoforms (i.e. post-translational modification^6^ or alternative splicing^7,8^) across biological conditions might result in differences in quantities for peptides that harbor those protein subregions. For example, if a protein quantity is unchanged, but one PTM site is increased with our treatment, that would cause the apparent disappearance of the peptide sequence that harbors that PTM (**Figure 1B**). Further, analytical problems during both sample preparation and data analysis can produce low quality peptide quantitation for a subset of peptides mapped to a protein, which would be desirable to remove^9^. Therefore, strategies are needed to assess concordance of quantitation for multiple peptides mapping to the same protein.

**Figure 1:**
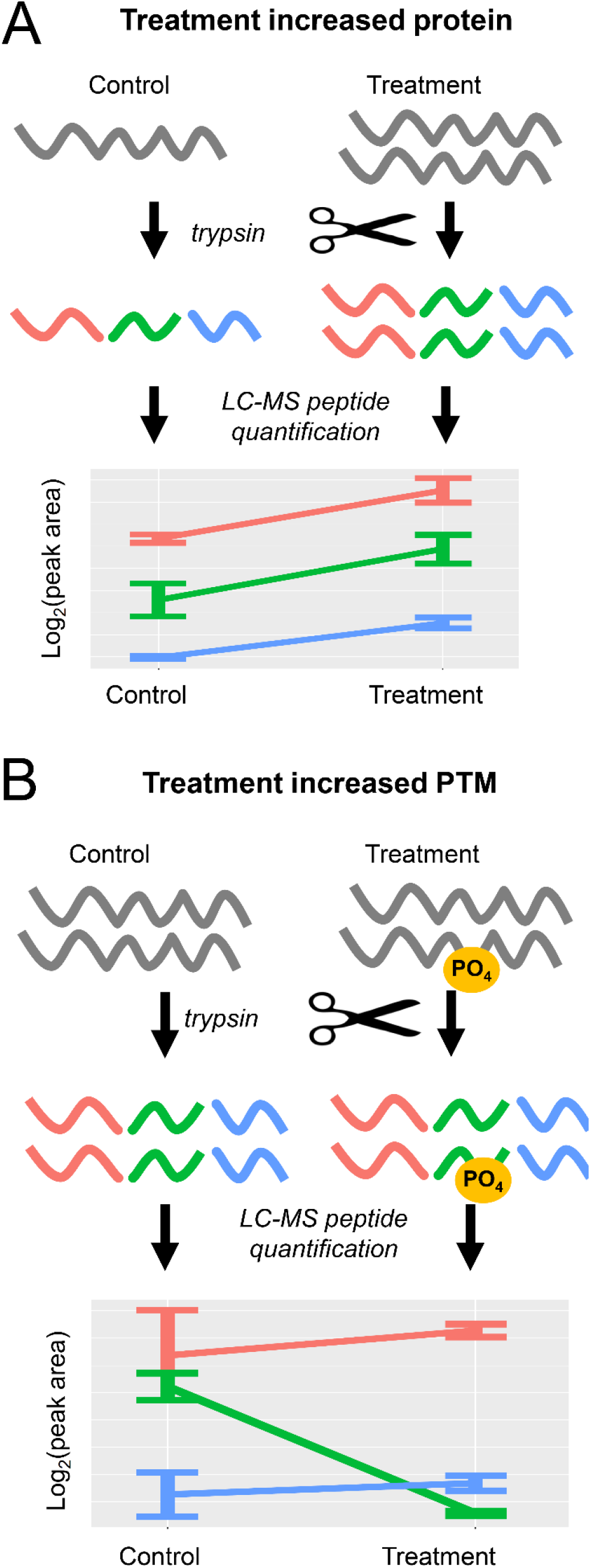
Protein quantification based on multiple measures of peptide parts. In shotgun proteomics, peptides are produced from proteins by proteases, and then the quantities of those peptides are used as a proxy for the protein. (A) Illustration of the case where a protein is increased by a biological treatment. The quantitative values from multiple peptides from one protein change in the same direction across biological treatment groups. (B) Illustration of the case where a protein modification is increased due to a biological treatment, but the protein quantity is unchanged. The quantitative values for multiple peptides from that one protein disagree.

This work presents a simple strategy called peptide correlation analysis (PeCorA) to detect peptides with discordant quantitation across treatment groups. PeCorA uses a linear model to statistically assess the interaction between peptide and treatment groups. Peptides with quantitative patterns that differ from all other peptides in that protein are reported as potentially interesting. Examples of those peptides include direct detection of a regulated methionine oxidation in PKA R1α, indirect detection of a lost phosphorylation in VAV1 protein, and incorrect peak picking of a peptide from CALR. PeCorA can be easily applied to any dataset and is freely available implemented in R.

## EXPERIMENTAL

### Data

Raw proteomic data from a published study of mouse primary microglia^10^ was downloaded from the Pride repository^11^ (identifier PXD014466, https://www.ebi.ac.uk/pride/archive/projects/PXD014466). The data set was composed of five biological replicates each from three sample groups: control, 50 mM ethanol treatment, or 5 ng lipopolysaccharide (LPS) treatment (15 total files). Each sample was analyzed with a 120-minute liquid chromatography gradient and online electrospray ionization into a hybrid quadrupole-orbitrap mass spectrometer (Q-Exactive Plus). Please see the original publication for more data collection and sample preparation details.

### Peptide Identification by database search of MS/MS spectra

All raw files from the microglia dataset were converted to mzML format using msconvertGUI (part of ProteoWizard)^12^ and searched against mouse proteins including isoforms downloaded from uniprot (04/08/20) with MS-Fragger version 2.4^13^. Reversed protein sequences and common contaminants were added to the database by the Philosopher toolkit version 2.0.0 (https://philosopher.nesvilab.org/). Searches were performed using the FragPipe user interface with the default closed search settings except 10 ppm precursor and 20 ppm fragment mass tolerances were used. Search outputs of each separate LC-MS analysis were refined with PeptideProphet^14^ and combined using iprophet^15^.

### Peptide quantification

Filtered peptide identifications were imported into Skyline for quantification^16^ by MS1 filtering^17^,^18^. Only protein identifications supported by two peptides were included, and peptides that were not unique to only one protein accession were excluded. Precursor signal within 10 ppm of the theoretical peptide mass for charge states 2-5 was extracted within 5 minutes of peptide identifications, and peaks were automatically picked by the software.

### Peptide Correlation Analysis (PeCorA)

Total peak area for each peptide precursor was exported from Skyline (report template PeCorA.skyr available with data repository) with the protein annotation, condition group, biological replicate number, and charge state. This report was read into R for processing. First, the values were filtered to include only precursors with measured MS1 areas in all samples. Next, the peak areas were log transformed, and the global distribution of all peak areas was scaled to have the same center. After global scaling, each peptide scaling was performed to center each peptide relative to the mean of the control group’s peak area. This twice-scaled data was then used for peptide correlation analysis. At a high level, the slope of each peptide’s quantity across experimental conditions was compared with the slope of all other peptides measured from that protein. This was achieved by iteratively setting each peptide to its own factor group against the factor group of all other peptides, and then fitting a linear model that includes a term for the interaction between the peptide groups. The p-value of significance for an interaction was recorded for each peptide group in each iteration of the loop before moving to set the next peptide as the out group. All p-values for the peptide interaction terms within one protein were then correction with the Benjamini-Hochberg procedure^19^.

### Data and code availability

Peptide correlation analysis code is written in R and available on github from https://github.com/jessegmeyerlab/PeCorA. The github repository also contains the Skyline report template (skyr), skyline report from microglia data used here, and PeCorA outputs from analysis of the microglia data used here. The raw mass spectrometry data from the microglia dataset is available from the Pride repository of the original publication (PXD014466), and also on massive.ucsd.edu (dataset MSV000085712). The skyline file used to generate peptide areas from microglia data is available on Panorama at: https://panoramaweb.org/HRvw2O.url.

## RESULTS

The first step in PeCorA is data preparation. The global distribution of data is scaled to have the same center (**Figure 2A**), and then each peptide is scaled to the center of the control condition (**Figure 2B**). The former is standard practice in large-scale omics experiments, and the latter is required for PeCorA because we must align all peptides to the same scale if we want to accurately test if each peptide is different than all other peptides. Peptide scaling removes the spread of peptide quantities that results from differences in ionization efficiencies, and results in a much tighter distribution of peptide quantities for each protein. After the data is scaled, the PeCorA algorithm iteratively compares each peptide to all other peptides in the same protein (pseudocode in **Figure 2C**). At a high level, the slope of each peptide quantity is compared with the slope of all other peptide quantities across biological conditions using a linear model that includes an interaction term between the treatment and peptide groups. This enables calculation of a p-value testing whether each peptide agrees quantitatively with the group of all other peptides in that protein. All p-values for all peptides are recorded, and then multiple hypothesis testing is applied to adjust the distribution. Comparing the interaction p-value with and without peptide scaling demonstrates the improved statistical power (**Figure 2D**). Peptide quantities from the same protein (Psmd14) are used in **Figure 2B** and **Figure 2D** to illustrate these concepts. Thus, PeCorA is an algorithm that determines statistically whether a peptide’s quantity across experimental treatments disagrees with other peptides assigned to the same protein.

**Figure 2:**
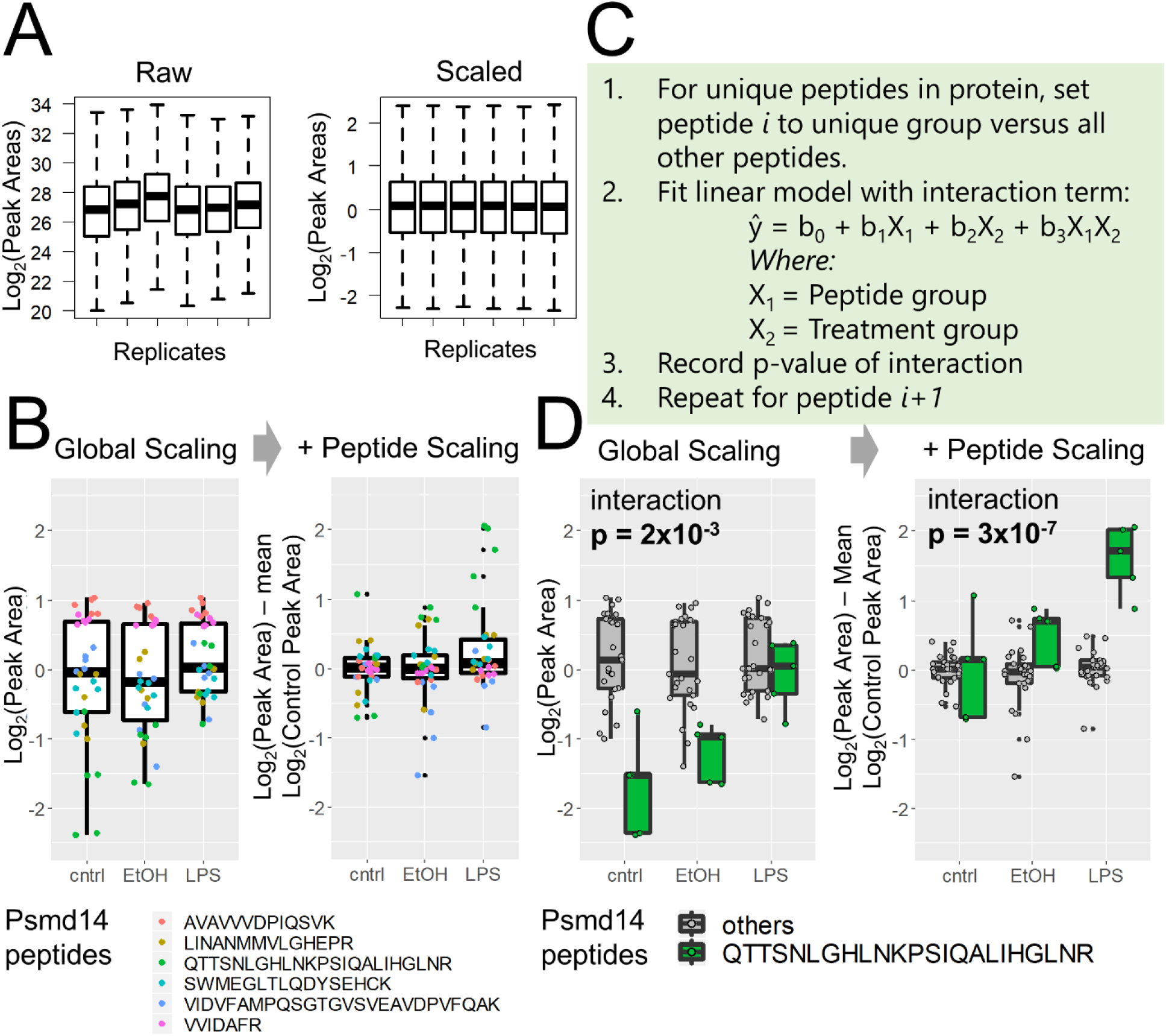
Peptide correlation analysis (PeCorA) to detect differentially modified peptides. (A) First, global data distributions are scaled to be centered around zero. (B) Next, each peptide is scaled to the mean of the control group to produce a unitless relative peptide quantity across groups. This helps correct the problem of differential peptide ionization and detection efficiency to produce more uniform data distributions. (C) Third, the quantitative values of one peptide from a protein is compared with the quantities of all other peptides in that protein using a linear model with a term for the interaction between peptides and biological treatment groups. This is repeated in a loop to compare each peptide to all other peptides. The p-value of the interaction is used to determine whether the quantity of each peptide is statistically different from all other peptides in that protein. (D) Example of one peptide that is statistically different from the quantities of all the other peptides. The p-value of the interaction between peptide and treatment group decreases when peptide scaling aligns the data.

The frequency of quantitative peptide disagreements in a dataset is useful first analysis to understand the potential utility of PeCorA. In total, after excluding peptides that map to multiple proteins, qualitative analysis of this dataset from microglia discovered a total of 27,685 peptides that map to a total of 2,918 proteins (requiring at least 2 peptides per protein). Data was further filtered to keep only peptides with peak areas over 100, and only peptides with quantities in all replicates, which left 26,444 peptides from 2,858 proteins. Of those, 416 proteins were found to harbor at least one peptide that disagreed with the other peptides in that protein, which corresponds to about 15% of quantified proteins. From the perspective of quantified peptides, the frequency of disagreement is much lower; 489 of the 26,444 peptides, or 1.9%, were found to disagree quantitatively with the other peptides mapped to the same protein. Overall, these results suggest that quantitative disagreements are rare at the peptide level, but relatively common from the protein perspective. Further investigation of these results reveal how PeCorA can discover multiple types of interesting peptide disagreements in a quantitative proteomics dataset: (1) direct evidence of regulated PTMs, (2) indirect evidence of regulated PTMs, and (3) incorrect peak picking.

As an example of direct evidence for a regulated PTM discovered by PeCorA, the peptide with the most significant adjusted p-value was from PKA R1α and contained methionine 331 oxidation (**Table S1, Figure 3A**). This peptide was detected with and without the oxidation, but only the oxidized peptide was increased due to LPS stimulation. PKA R1α is thought to be regulated by oxidative stress^20–22^, and the finding that LPS can induce this site-specific oxidation may prove to be yet another mechanism by which cellular metabolism is tuned. This site M331 is on the surface of second nucleotide binding domain (CNB-B) near the cAMP binding site^23^, and could regulate cAMP binding or protein-protein interactions (**Figure 3B**). This example of direct observation of this regulated PTM site uncovered by PeCorA would be missed using standard protein quantity summaries based on grouped peptide quantities.

**Figure 3 (previous page):**
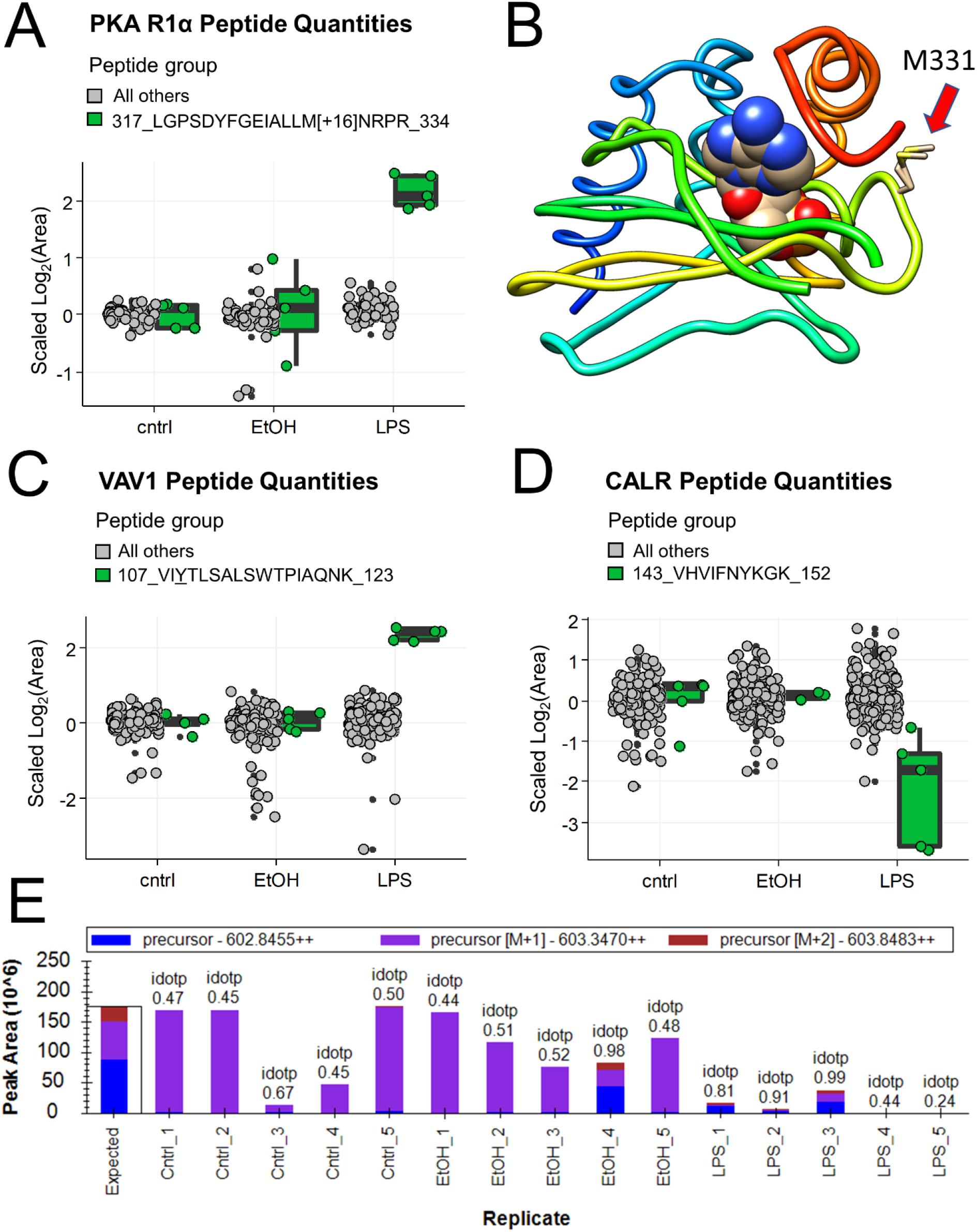
Examples of interesting peptides revealed by PeCorA. A. PKA R1α peptide quantities comparing the sequence containing oxidized methionine in green on the right with all other peptides in the same protein in grey on the left. B. Crystal structure (PDB: 5kjz) of the second nucleotide binding domain of PKA R1α bound to cGMP showing the location of the oxidized methionine 331 with a red arrow. C. VAV1 peptide quantities comparing the sequence with inferred change in phosphorylation in green on the right versus the all other peptides in grey on the left. D. CALR peptide quantities showing the peptide with problematic quantitation in green on the right with all other peptides in grey on the left. E. Peak area summary plot from Skyline for the peptide from CALR in (D) showing the poor isotopic dot product (idotp) reflecting incorrect peak picking across most

As an example of indirect evidence for an altered PTM, a peptide from the proto-oncogene Vav1 was found to increase due to LPS treatment (**Figure 3C**). Vav1 is known to signal downstream of tyrosine kinases, and this peptide revealed by PeCorA contained two known phosphorylation sites, pY110 and pS113. Prior literature provides strong evidence that this change in the unmodified peptide quantity may reflect a change in the abundance of the phosphorylated peptide form. Vav1 pY110 was decreased with IL-33 stimulation by 33%^24^, and IL-33 has been suggested as the signal mediating microglial response to LPS^25^. A decrease in the phosphorylation of this peptide would appear as an increase in the unmodified peptide quantified here. Further work is needed to verify if the peptide containing this phosphotyrosine is indeed altered in microglia in response to LPS stimulation, but the current evidence allows generation of hypothesis that unobserved PTMs are changing.

A third example shows the utility of PeCorA for quality control of peptide quantification. Due to the large number of peptides in any proteomics experiment, automated peak picking is required for quantification, which undoubtably leads to errors. In fact, active research is ongoing to develop methods that detect and exclude poorly quantified signals^9,26^. One peptide assigned to the protein Calreticulin (CALR) appeared to decrease due to LPS treatment (**Fig 3D**), but manual inspection of this peptide’s areas in Skyline revealed that the wrong peak was chosen for most replicates (**Fig 3E**). In summary, PeCorA was found to reveal poorly quantified peptides, and may find use as a filter to improve protein-level quantification in shotgun proteomics experiments.

## CONCLUSION

PeCorA is a new strategy to detect biologicially interesting protein sub-regions based on discordant peptide quantification across treatment groups. PeCorA is also useful for flagging peptides with poor quantitative reproducibility that should be excluded from analysis. PeCorA of proteomic data from mouse microglia treated with ethanol or LPS revealed that a significant proportion of proteins (15%) contain at least one peptide with discordant regulation. Such peptides may result from altered post-translational modification states, changes in protein sequence due to alternative splicing, or technical problems with peptide quantitation. PeCorA is an easy to use tool that should find widespread application to proteomic datasets.

## ACKNOWLEDGEMENTS

Thanks to the following people for sparking the idea of this paper via discussions on twitter: Karthik Kamath (@karthikskamath), Mike MacCoss (@mjmaccoss), Kareem Carr (@kareem_carr), Chelsea Parlett-Pelleriti (@ChelseaParlett), James Umbanhowar (@jumbanho), and Phil Wilmarth (@pwilmarth). Ronald Beavis (@NorSivaeb) helped find the microglia dataset.

## Notes

### Competing Interest Statement

The authors have declared no competing interest.

### Summary of Updates

Minor text updates, updated data availability

## References

1. Krey, J. F. et al. Accurate Label-Free Protein Quantitation with High- and Low-Resolution Mass Spectrometers. Journal of Proteome Research 13, 1034–1044 (2014).

2. Silva, J. C., Gorenstein, M. V., Li, G.-Z., Vissers, J. P. C. & Geromanos, S. J. Absolute Quantification of Proteins by LCMS^E^: A Virtue of Parallel ms Acquisition. Molecular & Cellular Proteomics 5, 144–156 (2006).

3. Tang, Liang. & Kebarle, Paul. Dependence of ion intensity in electrospray mass spectrometry on the concentration of the analytes in the electrosprayed solution. Analytical Chemistry 65, 3654–3668 (1993).

4. Cech, N. B. & Enke, C. G. Relating Electrospray Ionization Response to Nonpolar Character of Small Peptides. Analytical Chemistry 72, 2717–2723 (2000).

5. Aebersold, R. et al. How many human proteoforms are there? Nature Chemical Biology 14, 206–214 (2018).

6. Mair, W. et al. FLEXITau: Quantifying Post-translational Modifications of Tau Protein *in Vitro* and in Human Disease. Analytical Chemistry 88, 3704–3714 (2016).

7. Lau, E. et al. Splice-Junction-Based Mapping of Alternative Isoforms in the Human Proteome. Cell Reports 29, 3751–3765.e5 (2019).

8. Blencowe, B. J. The Relationship between Alternative Splicing and Proteomic Complexity. Trends in Biochemical Sciences 42, 407–408 (2017).

9. Tsai, T.-H. et al. Selection of features with consistent profiles improves relative protein quantification in mass spectrometry experiments. Molecular & Cellular Proteomics mcp.RA119.001792 (2020) doi:10.1074/mcp.RA119.001792.

10. Guergues, J., Wohlfahrt, J., Zhang, P., Liu, B. & Stevens, S. M. Deep proteome profiling reveals novel pathways associated with pro-inflammatory and alcohol-induced microglial activation phenotypes. J Proteomics 220, 103753 (2020).

11. Perez-Riverol, Y. et al. The PRIDE database and related tools and resources in 2019: improving support for quantification data. Nucleic Acids Research 47, D442–D450 (2019).

12. Kessner, D., Chambers, M., Burke, R., Agus, D. & Mallick, P. ProteoWizard: open source software for rapid proteomics tools development. Bioinformatics 24, 2534–2536 (2008).

13. Kong, A. T., Leprevost, F. V., Avtonomov, D. M., Mellacheruvu, D. & Nesvizhskii, A. I. MSFragger: ultrafast and comprehensive peptide identification in mass spectrometry–based proteomics. Nature Methods 14, 513–520 (2017).

14. Keller, A., Nesvizhskii, A. I., Kolker, E. & Aebersold, R. Empirical Statistical Model To Estimate the Accuracy of Peptide Identifications Made by MS/MS and Database Search. Analytical Chemistry 74, 5383–5392 (2002).

15. Shteynberg, D. et al. iProphet: Multi-level Integrative Analysis of Shotgun Proteomic Data Improves Peptide and Protein Identification Rates and Error Estimates. Molecular & Cellular Proteomics 10, (2011).

16. MacLean, B. et al. Skyline: an open source document editor for creating and analyzing targeted proteomics experiments. Bioinformatics 26, 966–968 (2010).

17. Schilling, B. et al. Platform-independent and Label-free Quantitation of Proteomic Data Using MS1 Extracted Ion Chromatograms in Skyline: APPLICATION TO PROTEIN ACETYLATION AND PHOSPHORYLATION. Molecular & Cellular Proteomics 11, 202–214 (2012).

18. Meyer, J. Fast Proteome Identification and Quantification from Data-Dependent Acquisition–Tandem Mass Spectrometry (DDA MS/MS) Using Free Software Tools. Methods and Protocols 2, 8 (2019).

19. Benjamini, Y. & Hochberg, Y. Controlling the False Discovery Rate: A Practical and Powerful Approach to Multiple Testing. Journal of the Royal Statistical Society. Series B (Methodological) 57, 289–300 (1995).

20. Haushalter, K. J. et al. Cardiac ischemia-reperfusion injury induces ROS-dependent loss of PKA regulatory subunit RIα. American Journal of Physiology-Heart and Circulatory Physiology 317, H1231–H1242 (2019).

21. Brennan, J. P. et al. Oxidant-induced Activation of Type I Protein Kinase A Is Mediated by RI Subunit Interprotein Disulfide Bond Formation. Journal of Biological Chemistry 281, 21827–21836 (2006).

22. Srinivasan, S. et al. Oxidative Stress Induced Mitochondrial Protein Kinase A Mediates Cytochrome C Oxidase Dysfunction. PLoS ONE 8, e77129 (2013).

23. Lorenz, R. et al. Mutations of PKA cyclic nucleotide-binding domains reveal novel aspects of cyclic nucleotide selectivity. Biochemical Journal 474, 2389–2403 (2017).

24. Pinto, S. M. et al. Quantitative phosphoproteomic analysis of IL-33-mediated signaling. PROTEOMICS 15, 532–544 (2015).

25. Cao, K. et al. IL-33/ST2 plays a critical role in endothelial cell activation and microglia-mediated neuroinflammation modulation. Journal of Neuroinflammation 15, (2018).

26. Teo, G. et al. mapDIA: Preprocessing and statistical analysis of quantitative proteomics data from data independent acquisition mass spectrometry. Journal of Proteomics 129, 108–120 (2015).

